# Phytochrome gene expression studies: PHYE is required for FR-induced expression of PHYA and PHYD suppresses expression of PHYA

**DOI:** 10.1101/2021.06.20.449137

**Authors:** Umidjon Shapulatov, Mara Meisenburg, Mark van Hoogdalem, Alexander van Hall, Wim van Ieperen, Maarten van Wassenaar, Alexander van der Krol

**Affiliations:** Laboratory of Plant Physiology, Wageningen University & Research, Droevendaalsesteeg 1, 6708 PB Wageningen, The Netherlands; Horticulture and Product Physiology, Department of Plant Sciences, Wageningen University and Research, Droevendaalsesteeg 1, 6708 PB Wageningen, The Netherlands; Faculty of Biology, National University of Uzbekistan, University street 4, 100174 Tashkent, Uzbekistan; Business Unit Greenhouse Horticulture, Wageningen University & Research, Droevendaalsesteeg 1, 6708PB Wageningen, The Netherlands

**Keywords:** PHYTOCHROME, LED, Luciferase reporter plants, transcription

## Abstract

Arabidopsis has five phytochrome (PHY) genes for sensing the Red:Far Red (R:FR) ratio in ambient light, of which PHYA has an established role in responses to FR. To study whether and how PHYs may influence each other’s transcription, PHY-Luciferase reporter plants (pPHYA:LUC, pPHYB:LUC, pPHYC:LUC, pPHYD:LUC and pPHYE:LUC) were constructed. Subsequently, reporter lines representative for each PHY were crossed into each of the five single *phy-*mutant backgrounds. Reporter activities in WT and *phy* mutant was studied under diurnal mixed (R, B, FR), R, FR or B LED light in seedling or rosette plants. Both pPHYA:LUC and pPHYB:LUC show strong induction under FR. Full FR upregulation of both PHYA and PHYB is dependent on PHYE, identifying PHYE as a novel sensor for FR light responses. Results also show that PHYA expression is strongly suppressed by PHYD. Results were confirmed for expression of endogenous PHYA and PHYB, albeit with different dynamics compared to the LUC reporters. Profiling of pPHYA:LUC and pPHYB:LUC reporters suggest gating of FR responses. Manipulation of PHY expression levels by FR may provide a novel basis for manipulating plant growth in controlled environments.

## Introduction

The most important photoreceptors that control plant growth as function of the Red (R) and Far-Red (FR) light spectrum are a family of phytochrome (PHY) genes, which in Arabidopsis consist of PHYA-PHYE (Bae and Choi 2008). Phytochromes are produced in the inactive red (R) light absorbing Pr form. Upon perception of red light the inactive Pr state of phytochromes changes to the active Pfr state to trigger responses in both the cytosol (Paik, Yang et al. 2012) and in the nucleus (Nagy and Schafer 2002, Nagatani 2004, Kevei, Schafer et al. 2007, Van Buskirk, Decker et al. 2012, Klose, Viczian et al. 2015). In the nucleus phytochrome protein interacts with multiple Phytochrome Interacting Factors (PIFs) to mediate light transcriptional responses (Huq, Al-Sady et al. 2004, Castillon, Shen et al. 2007, Leivar and Quail 2011). While phytochromes are activators, Phytochrome Interacting Factors (PIFs) are considered repressors of photomorphogenesis, because the interaction of phytochrome Pfr with PIFs promotes their turnover (Park, Park et al. 2012, Xu, Paik et al. 2015). The interaction between phytochromes and PIFs do not only result in degradation of the PIFs, but also in co-degradation of the phytochrome protein. Indeed, the function of Pfr in the nucleus is controlled by multiple nuclear factors that are involved in nuclear Pfr stability (Monte, Tepperman et al. 2004, Khanna, Shen et al. 2007, Al-Sady, Kikis et al. 2008, Leivar, Monte et al. 2008, Leivar and Quail 2011, Ni, Xu et al. 2013). It has been shown that PIFs regulate phyB-E protein stability through COP1/DET/FUS (Jang, Henriques et al. 2010). In addition, PIFs and PHYs interact with a CUL3-based E3 ubiquitin ligases complex containing the Bric-a-Brac/Tramtrack/Broad Complex (BTB)-domain containing substrate adaptor Light-Response (LRB). Presumably PIFs and PHY are co-degraded by interaction between a CUL3-LRB-PIF complex and a CUL3-LRB-PHY complex, through dimerization of the LRBs (Christians, Gingerich et al. 2012). Translocation of PHY proteins into nucleus is required for the nuclear signaling and the translocation of PHYA Pfr protein into the nucleus is controlled by the FAR-RED ELONGATED HYPOCOTYL 1 (FHY1) AND FHY1-LIKE (FHL) (Genoud, Schweizer et al. 2008). PHYs also have a function in the cytosol where they control translation of specific mRNAs (Paik, Yang et al. 2012). The stability of the pool of cytosolic Pfr is regulated by cytosolic factors, explaining why the dynamics of nuclear PIF protein turnover and total PHY protein turnover may not be the same.

Transcription of PHY genes and translation of the PHY mRNAs determine the actual pool of phytochrome protein that is available for PHY protein signaling. Understanding transcriptional regulation of phytochrome genes is therefore integral part of understanding overall phytochrome action. Phytochrome genes are regulated by the circadian clock (Toth, Kevei et al. 2001), while in turn the clock is entrained through phytochrome signaling (Somers, Devlin et al. 1998). Genetic interactions between phytochromes affect germination, hypocotyl elongation and flowering, but this was not correlated to possible effects on PHY gene transcription (Sanchez-Lamas, Lorenzo et al. 2016). Recently it was shown that the promoter of PHYA is targeted by PIF4 and PIF5 (Seaton, Toledo-Ortiz et al. 2018), which potentially couples PHYA transcription to R:FR light conditions, as R:FR conditions determine the interaction between PHYB and PIF4/5 and the stability of these proteins (Lorrain, Allen et al. 2008, Foreman, Johansson et al. 2011).

In the studies on phytochrome action the effects of a given light treatment on PHY promoter activities are usually ignored. However, whether this is always justified under day light conditions or under artificial LED light with its unnatural spectral composition of LED lights requires verification. Indeed, there is no comprehensive and systematic analysis of PHY gene transcription as function of (LED) light quality. Therefore, we investigated PHY promoter activity as function of different LED light conditions at different developmental stages and as function of individual phytochromes. Dynamic transcriptional responses *in planta* was monitored using firefly luciferase (LUC) reporter lines (Millar, Short et al. 1992). The pPHYA:LUC, pPHYB:LUC, pPHYC:LUC, pPHYD:LUC and pPHYE:LUC reporter lines were crossed into each of the single phytochrome mutant backgrounds, resulting in a total of 30 reporter lines. Results show that interactions between phytochromes at the transcription level change from seedling to mature rosette stage. The diurnal pPHY:LUC activity was monitored in response to day-time R, FR or B LED light, showing strong upregulation of pPHYA-LUC and pPHYB-LUC activity under FR. Moreover, this induction by FR was not dependent on the classical FR light sensor PHYA. Full FR-induction of PHYA and PHYB under FR depends on PHYE, thus identifying PHYE as novel sensor for FR. Similarly, PHYE affected expression of endogenous PHYA and PHYB under FR. The studies thus show unexpected complex PHY-interaction in the regulation of PHY gene expression.

## Materials and methods

### Plant materials and growth conditions

Seeds of *Arabidopsis thaliana* T-DNA insertional mutant lines were obtained from the Nottingham Arabidopsis Stock Centre (NASC, University of Nottingham, UK). The following lines were used in our work: WT (Col-0), ph½4-T(NASC: N661576), *phyB-9* (Reed, Nagpal et al. 1993), *phyC-2* (N66036), *phyD* (N676270), *phyE-T* (N671700). All phytochrome mutants are in Col-0 background. The *phy* T-DNA insertion mutants were validated as homozygous insertion mutant by PCR of genomic DNA using Salk T-DNA and gene specific primers (Table S1). For Luminator (van Hoogdalem, Shapulatov et al. 2021) experiments, seeds were sawn on MS-0.8% agar plates (Murashige-Skoog medium 0.22g/L, 8g/L plant agar Duchefa), stratified in the dark for three days at 5°C, after which they were sown on 4×4×4cm rockwool blocks (Grodan, Roermond, The Netherlands) soaked in Hyponex nutrient solution (Unifarm, Wageningen, The Netherlands). Plants were pre-grown in a climate chamber (12hL/12hD; 22°C; relative humidity (RH) at 65%). Directly before transfer to LUMINATOR, reporter plants were watered by soaking the rockwool blocks in Hyponex solution, which allows for growth for up to 6 days without additional watering. Light conditions in LUMINATOR cabinet are described below.

### PHY-LUC reporter cloning and construction homozygous reporter lines

Construction of the pPHY:LUC reporter genes using ~2kb upstream promoter fragments of either PHYA, B, C, D or PHYE is described in (Toth, Kevei et al. 2001). Binairy vectors containing these reporter genes were kindly donated by the group of Prof. Nagy.

Arabidopsis Col-0 plants were transformed by floral dip transformation (Zhang, Henriques et al. 2006) and positive transformants were selected based on Luc activity. Representative homozygous lines were crossed into the different phytochrome mutant backgrounds. For all progeny homozygous for the pPHY:LUC reporter, the homozygous *phy*-mutant genotype was confirmed by PCR and T_4_ plant homozygous for both the phytochrome mutation and the respective pPHY:LUC reporter were used.

### In planta LUC reporter activity measurements in LUMINATOR

LUC activity in the different pPHY:LUC reporter plants was measured in a custom built LUMINATOR cabinet containing a high performance PIXIS: 1024 CCD camera (Princeton Instruments, Roper technologies, Sarasote, FL, USA) fitted with a 35mm f/1.4 Nikkor SLR lens (Nikon, Shinjuku, Tokyo, Japan). Plants were pre-sprayed with 1mM D-luciferin (Promega, Fitchburg, WI, USA) one day before imaging to deplete accumulated LUC (de Ruijter, Verhees et al. 2003). For multiple day measurements plants were sprayed daily with 1 mM D-luciferin (Promega, Fitchburg, WI, USA) at 10 am. Plants were acclimated to conditions in LUMINATOR for one day. LUC activity images were taken every 30 minutes with an exposure time of 7 minutes, during which LED illumination is switched off. Light from chlorophyll fluorescence of plants was blocked by using a ZBPB074 Bandpass Filter (Asahi Spectra, Sumida, Tokyo, Japan).

### Light conditions during diurnal LUC activity imaging

Imaging of ff-LUC activity in plants is under mixed LEDs, emitting R (590-660nm), B (420-500nm) and FR 680-760). In addition, a ramping in R intensity and R:FR ratio was used to mimic natural morning and evening light light conditions. The light intensity during 2 hours ramping at start-day and end-day is 33 μmole m^-2^ s^-1^and during the remaining hours of the photoperiod 90 μmole m^-2^ s^-1^. Photosynthetically active radiation (PAR) intensity was 25 and 80 μmole m^-2^ s^-1^ respectively. The ratio B:R:FR light during ramping is 1:2:1 and during the remaining hours of the photoperiod 3:6:1.

The Red light treatments were at 80 μmole m^-2^ s^-1^ of pure Red light, the FR light treatment was at 430 μmole m^-2^ s^-1^ of FR LED light and the Blue light treatment was at 30 μmole m^-2^ s^-1^ of blue light. The R>FR step gradient light treatment consists of 3 hours R:FR=8, 3 hours R:FR=1 (mild shade), 3 hours R:FR=0.5(shade) and 3 hours R:FR=0.2 (deep shade). PAR intensity was 80-85 μmole m^-2^ s^-1^ during all shade conditions. Light quality/intensity was measured the using Flame-T spectroradiometer (Ocean Optics, Duiven, The Netherlands). Relative luminescence was quantified in Image J (imagej.nih.gov/ij) as described before (Shapulatov, van Hoogdalem et al. 2018, van Hoogdalem, Shapulatov et al. 2021).

### Quantitative RT-PCR

Total RNA was isolated from 3 weeks old rosettes that placed under mixed LED for 1 day for acclimatization and the next the day, plants were exposed to far-red light (430 umol /m^2^s). rosettes were harvested at the indicated times. mRNA and cDNA synthesis were as described (van Hoogdalem, Shapulatov et al. 2021). qRT-PCR for quantification of endogenous PHY genes on a CFX Connect Real-time system machine (BioRad, CA, USA). Primer sequences used can be found in Table S1. Statistical analysis of the qPCR data was carried out in RStudio (3.6.0). A 2-way ANOVA was used to analyse the qPCR data followed by a Least Significant Different (LSD) post-hoc test (agricolae 1.3-5, R package). The datasets for the PHYA and PHYB activity were first log transformed to meet the assumptions of the 2-way ANOVA.

## Results

### Construction of thirty pPHY:LUC reporter lines

In order to study the expression of the five phytochrome genes (PHYs) of Arabidopsis in WT and phytochrome mutant backgrounds, the upstream 2-2.5 kb promoter of each of the five PHY genes was fused to the firefly luciferase (LUC) coding sequence in binary expression vectors (Toth, Kevei et al. 2001), resulting in five pPHY:LUC reporter constructs. The different pPHY:LUC reporter constructs were introduced into Arabidopsis WT (Col-0) by the agrobacterium mediated floral dip transformation (Zhang, Henriques et al. 2006). For each of the pPHY:LUC reporters a minimum of ten primary transformants were screened for luciferase activity and one representative transformed plant was selected and developed into a homozygous reporter line expressing either pPHYA:LUC, pPHYB:LUC, pPHYC:LUC, pPHYD:LUC or pPHYE:LUC. Subsequently, each of the five homozygous pPHY:LUC reporter plants was crossed to each of five single *phy*-mutant plants. Phytochrome mutant backgrounds were selected based on seedling growth phenotype under specific light conditions and PCR analysis of genomic DNA using specific primers (Table S1) (Nagatani, Reed et al. 1993, Hennig, Funk et al. 1999, Balasubramanian, Sureshkumar et al. 2006, Chen, Sonobe et al. 2013). By crossing the pPHY:LUC reporter into the different phy-mutant backgrounds, the relative expression of the LUC reporter in WT and mutant can be compared directly, since the reporter is in same chromosomal location with identical ‘position’ effects on transgene expression. The pPHY:LUC reporter lines and constructs are listed in Table S2. Fig. 1 shows representative images of the pPHY:LUC reporter activity in WT in three week old rosette plants. The relative level of LUC activity is not the same for the different PHY reporters. At the rosette stage the PHYA and PHYC promoters show the strongest transcriptional activity, while transcription from the PHYE promoter is very weak.

**Fig.1.**
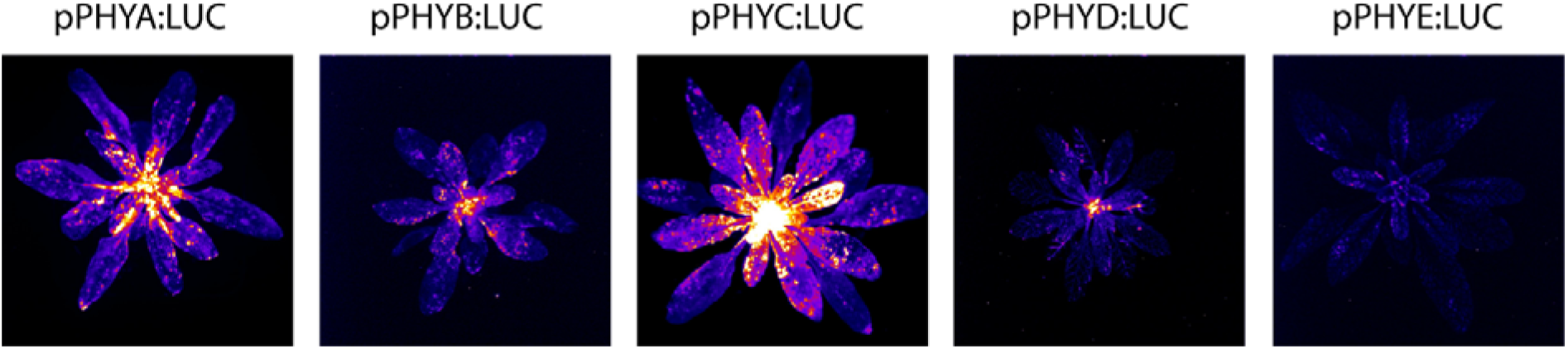
pPHY:LUC reporter plants. Relative luciferase activity was captured in 28-day old rosette plants sprayed with 1 mM luciferin-D. LUC activity image capturing was by seven minutes exposure time. pPHYC:LUC shows the highest activity, while pPHYE:LUC is a little above background.

### PHY mediated genetic interactions on PHY-LUC activity diminish during plant maturation

pPHY:LUC activity in WT and *phy* mutants was imaged in 7 and 14 day old seedlings and 24-26 day old plants. For all three developmental stages the LUC activity imaging was at 11 am, close to the maximum phase of pPHY:LUC reporters in seedlings (Toth, Kevei et al. 2001). The average relative LUC activity per seedling/plant was quantified for each of the reporter lines (Fig. 2). The results indicate that especially in 7-day-old seedlings for many pPHY-LYC reporters the activity is significantly increased in a *phy*-mutant background, indicating that at early stages of development different phytochromes are involved in repression of transcription of the pPHY-LUC reporters (Fig. 2A-E). However, as plants mature, this genetic interaction between phytochromes at the transcription level diminishes (Fig. 2A-E). Most remarkable is the consistent elevated level of pPHYA-LUC expression in the *phyD* mutant background, indicating that PHYD is a constitutive suppressor of pPHYA-LUC transcription. As plant mature the genetic interactions between the different phytochromes diminishes, but is still apparent for the suppression by PHYB on pPHYA-LUC, the suppression by PHYB, PHYC, PHYD and PHYE on pPHYB-LUC and suppression by PHYA, PHYB and PHYC on pPHYD-LUC (Fig. 2, 24 day old plants). In contrast, full expression of pPHYC-LUC in mature plants requires PHYB (Fig. 2C, 14 and 24 day old plants).

**Fig.2.**
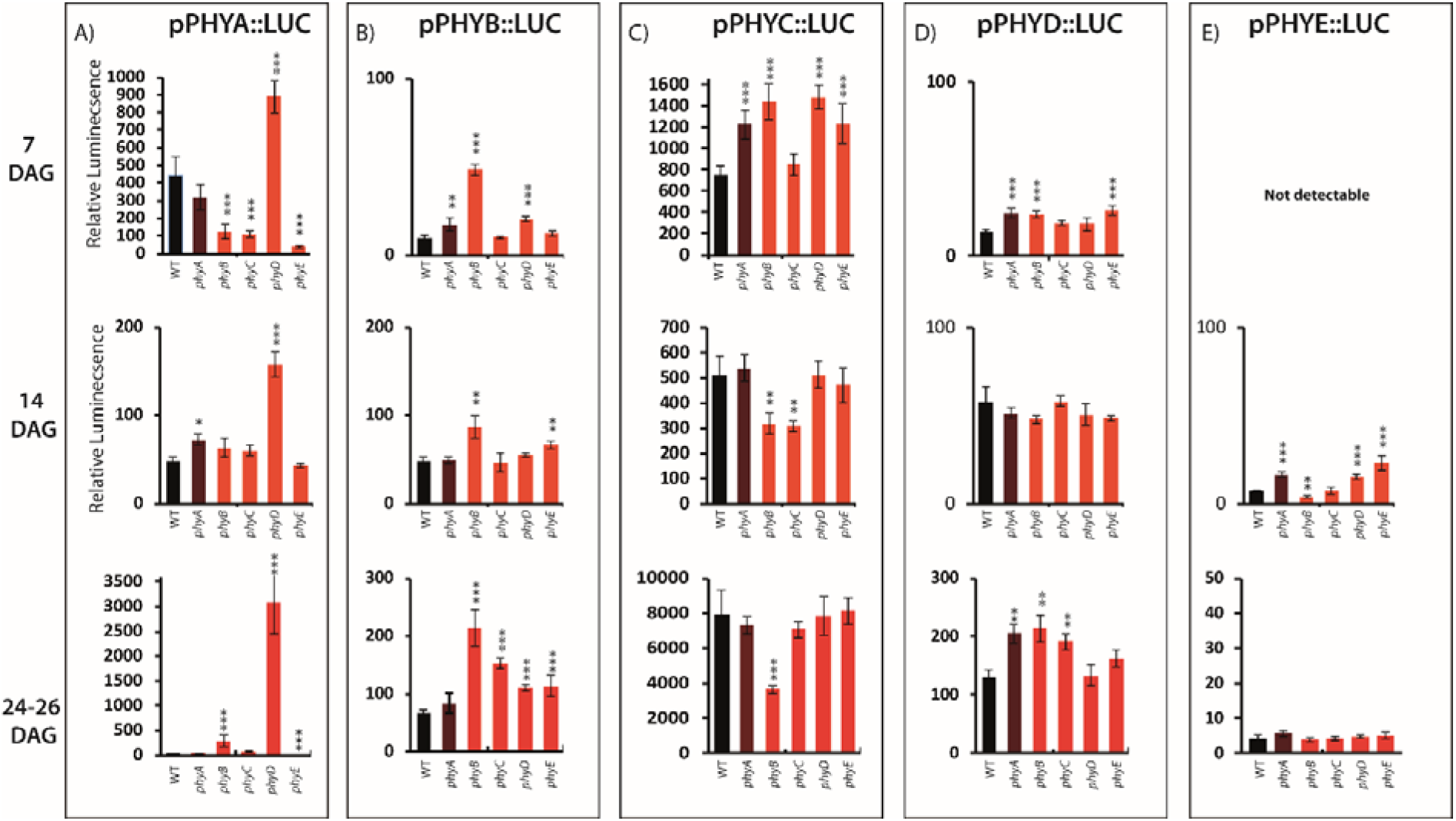
pPHY:LUC activity at ZT=3hr at different developmental stages. Plants were grown in growth cabinets under fluorescent WL and sprayed with substrate luciferin (1 mM) one day and one hour before imaging. LUC activity was imaged in plants at 7, 14 and 24-26 days post germination at 11 am (ZT=3hr). A: pPHYA:LUC in WT and five phytochrome mutants; B: pPHYB:LUC in WT and five phytochrome mutants; C: pPHYC:LUC in WT and five phytochrome mutants; D: pPHYD:LUC in WT and five phytochrome mutants; E: pPHYE:LUC in WT and five phytochrome mutants (not detectable at 7 days). The relative LUC activity was quantified in ImageJ. Number of replicate plants for each reporter line: N=9 for 7 DAG, N=9 for 14 DAG and N=6 for rosette plants. Error bars represent mean ±SE. Error Bars with symbols (*; **; ***) indicate a significance to compare WT respective to p-value <0.05; <0.01; <0.001.

### pPHY-LUC responses to R, B and FR light

To determine the pPHY:LUC activity in WT under different light conditions, plants were imaged in Luminator (van Hoogdalem, Shapulatov et al. 2021) for four days. After one day acclimation, LUC activity was imaged in plants under 12hrR/12D, followed by 12hrFR/12D and finally 12hrB/12D, using either R, B or FR LED light during the day. The pPHY:LUC activity images were obtained every 30 minutes. This was done for 7-day old plants, 14-days old plants and when plants were 25-day old. Qualitatively the responses of the different pPHY:LUC reporters are similar at these three stages of development and results of the expression profiles in 14 day old plants are shown in Figure 3. Most pPHY:LUC reporters do not show a strong response to the R photoperiod, except for pPHYC:LUC which is induced under R. Also, pPHYC:LUC shows a consistent transient increase in activity at the day-night transition following all photoperiods (Fig. 3). Most remarkable is the strong and immediate upregulation of pPHYB:LUC under FR light following the dark period of the night, reaching a peak expression almost 10-fold higher then under R light. During the night following FR, both expression of pPHYB:LUC and pPHYA:LUC show an initial rapid decline. Expression of pPHYA:LUC is also upregulated by FR light but in a more gradual way, reaching a 6-fold higher expression at the end of the FR photoperiod compared to under R. Moreover, only pPHYC:LUC shows a transient increase in activity at the day-night transition following all photoperiods. The other pPHY:LUC reporters were not induced by FR, or showed a decline of expression under FR. Only pPHYB-LUC and pPHYC-LUC show a strong response to B during the day (Fig. 3). Overall, results show that light quality may change the expression level of some of the phytochrome genes and thus light quality changes the potential pool of phytochrome protein that may participate in light signaling. Note that some of the fine structure in the LUC activity profiles during the day is most likely due to renewed spraying of luciferin and some leaf hyponastic movement.

**Fig.3.**
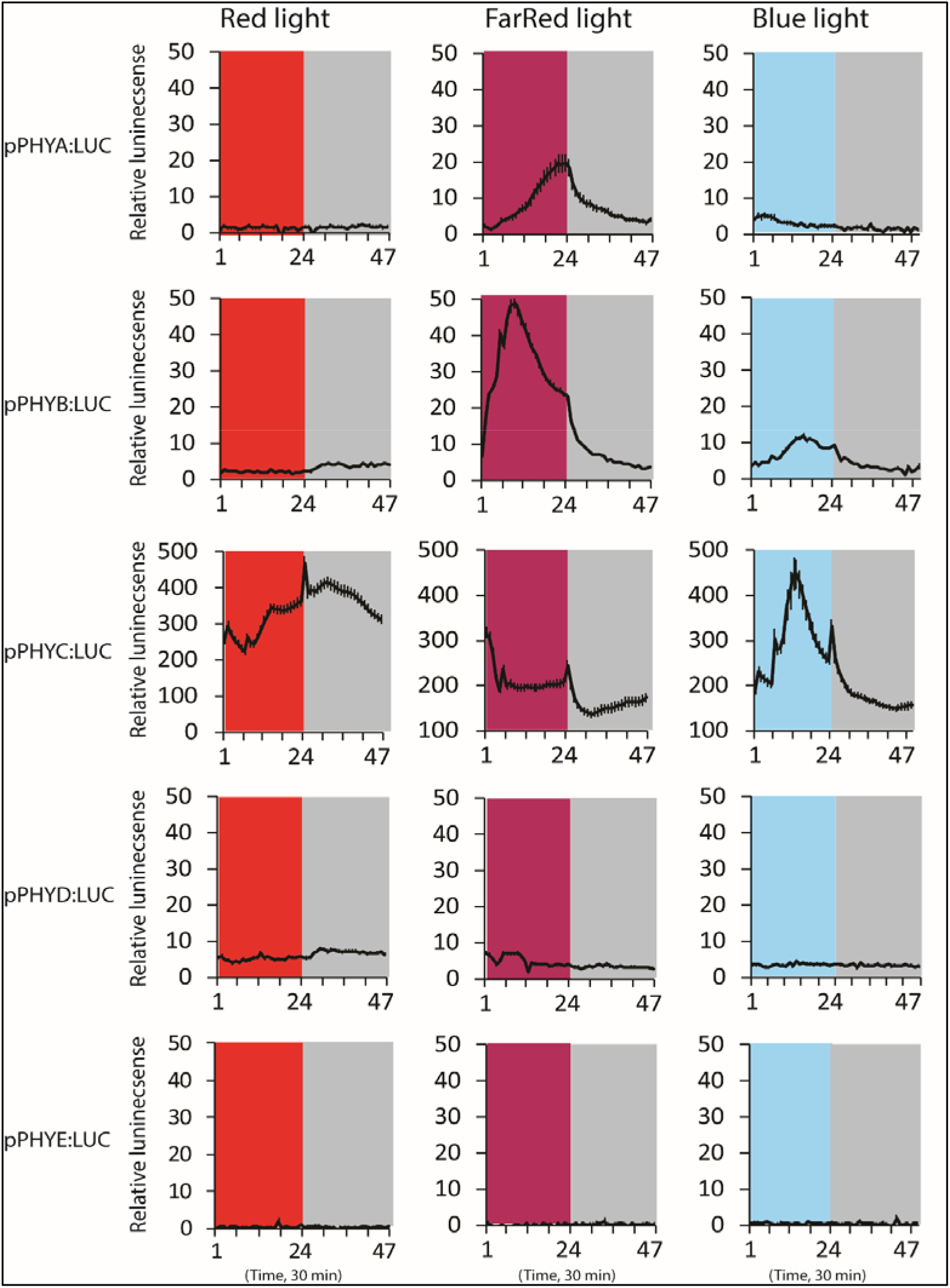
Diurnal expression profile of the pPHY:LUC reporters under diurnal 12R/12D, 12FR/12D and 12B/12D. Plants were grown under 12L/12D and 14 days after germination seedlings were sprayed with 1 mM luciferin. One day later plants were placed in LUMINATOR for adjustment under diurnal R+B+FR for one day. Subsequently plants were exposed to light regimes 12R/12D, followed by 12FR/D and finally 12B/12D. Luciferin (1 mM) solution was sprayed once per day. LUC activity images were obtained every half hour (7 min. exposure) for each full diurnal cycle. The relative LUC activity was quantified in ImageJ and corrected for background signal. Number of replicate seedlings for each reporter line: N=6. Error bars represent mean ±SE.

### Feedback regulation on pPHYB-LUC by PHYB and PHYE under FR and B

Because phytochromes are involved in R:FR sensing, the strong induction of pPHYB:LUC in WT plants under FR light suggests the involvement of PHYA, which is the classical sensor for FR light responses (Whitelam, Johnson et al. 1993, Yanovsky, Casal et al. 1997, Fankhauser 2001). To determine the role of individual phytochromes in expression of pPHYB:LUC under different light conditions (R/FR/B, R, FR or B LED light), the pPHYB:LUC activity was monitored in the different phytochrome mutant backgrounds (Fig. S1). As example, here we show results for pPHYB:LUC in WT and different phytochrome mutants under FR and B (Figure 4). The induction of pPHYB:LUC under FR is significantly reduced in the *phyB* and the *phyE* mutant background, indicating that in context of transcriptional regulation of pPHYB:LUC expression the PHYB and PHYE act as a FR sensor. In contrast, the classical FR sensor PHYA has little effect on pPHYB:LUC activity under FR (Fig. 4). Most phytochromes do not affect the response of pPHYB:LUC to B, except for PHYC, which represses pPHYB:LUC activity in WT (Fig. 4).

**Fig.4.**
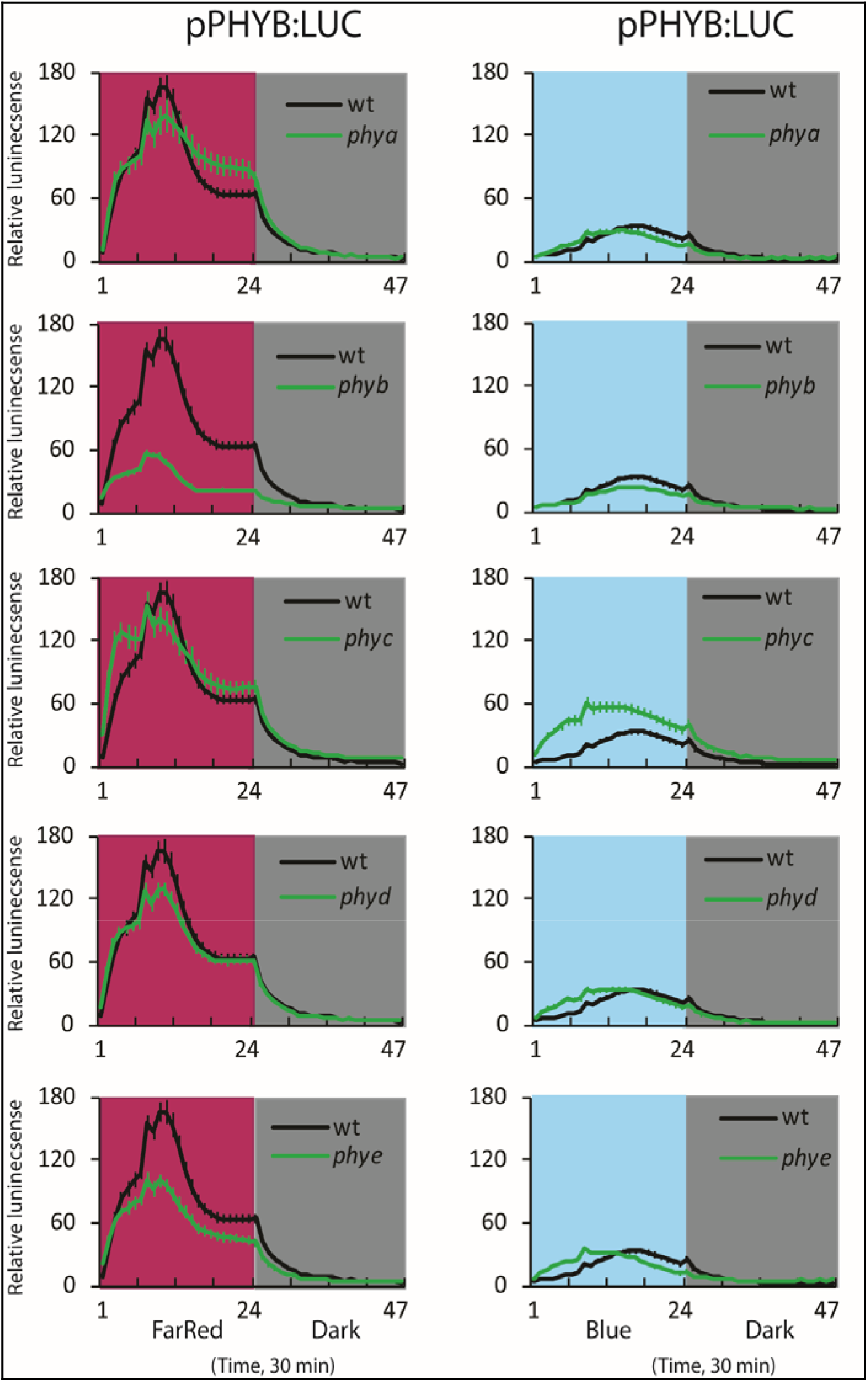
pPHYB:LUC activity in WT and phytochrome mutants in 14 day old seedlings under FR or B. Seeds of the pPHYB:LUC reporter lines were stratified and germinated in growth cabinets under diurnal fluorescent WL (12L/12D). At 14 days after germination seedlings were sprayed with substrate luciferin (1 mM) and one day later placed in LUMINATOR for adjustment under diurnal R+B+FR for one day. Then plants were exposed to light regimes of 12mixed/12D, 12R/12D, 12FR/D and finally 12B/12D. Luciferin (1 mM) solution was sprayed once per day. LUC activity images were obtained every half hour (7 min. exposure) for each full diurnal cycle. The relative LUC activity is quantified in ImageJ and adjusted for background signal. Number of replicate seedlings for each reporter line: N=16. Error bars represent mean ±SE. All results are shown in Fig. S3. Here only results for pPHYB-LUC in WT and *phy*- mutants under FR and B are shown.

The diurnal pattern of PHYB promoter activity under the different light regimes indicate that the phase of pPHYB:LUC activity is dependent on the light conditions (phase of pPHYB:LUC in WT under mixed LED ZT=3 hr, under R ZT=4 hr, under FR ZT= 6 hr; Fig. S1). In addition, the phase is dependent on the phytochrome mutant background (phase of pPHYB:LUC under B in WT ZT=3 hr, in *phyB* mutant ZT=7 hr, in *phyC ZT*= 2 hr; Fig. 4). Expression of pPHYB:LUC in the *phyB* mutant background under mixed or R light is increased compared to in WT, but decreased under FR and B LED light compared to in WT (Fig. S1). This indicates that the effect of PHYB on its own promoter activity is dependent on light conditions and may switch from a repressor interaction (under mixed and R light) to activator interaction (under FR and B) (Fig. S1 and Fig. 4).

### pPHY-LUC response to “shade” depends on time of the day

To investigate the phytochrome gene expression as function of shade light conditions, pPHY:LUC reporter activity was imaged in 25-day old WT rosette plants placed under varying ratio’s of R:FR light. The five sets of WT pPHY:LUC reporter plants were placed in LUMINATOR to adapt for two days to diurnal mixed LED light (R,B,FR). After the night of the second day, the photoperiod was started using 3 hours of R light combined with low level of FR (R:FR=8). Subsequently, every 3 hours the R level remained the same, but dosage of FR was increased going from R:FR=8 to R:FR=1, to R:FR=0.5 and finally ending the day with 3 hours of R:FR=0.2, which mimics deep shade conditions. After the night following these 4 blocks of increasing shade light conditions, the next day, the same blocks of R+FR LED light were given in reverse order, starting the day with R:FR=0.2 and ending the day with R:FR=8. The different pPHY:LUC reporters show different responses to the shade treatments (Fig. 5). Both pPHYA:LUC and pPHYB:LUC show little response to the day under mixed R,FR,B LED and no response to 3 hr R:FR=8 (Fig. 5). Under R:FR=1 pPHYB:LUC shows an direct transcriptional response, while for pPHYA:LUC and pPHYC:LUC a transcriptional response only starts near the end of three hour R:FR=1 (Fig. 5). In contrast pPHYD:LUC expression is down regulated during this light treatment. This is consistent with our discovery that PHYD is a suppressor of PHYA (Fig. 2) and suggests that part of the upregulation of PHYA may be caused by downregulation of PHYD under increasing shade conditions. However, the following day when light treatments are given in reverse order, pPHYD:LUC activity shows an increase at the end of 3hr R:FR=0.2, which is not mirrored by a decline in pPHYA:LUC activity (Fig. 5). During the night, expression of PHYA, PHYB and PHYC initially decline with different rates, which is less rapid than the decline in pPHY:LUC expression after pure FR (Fig. 3). Also, after pure FR expression declines to “normal” levels as seen under WL, mixed LED or B (Fig. 3), while after the R+FR light treatments the expression of pPHYA:LUC and pPHYB:LUC remains high throughout the night (Fig. 5).

**Fig.5.**
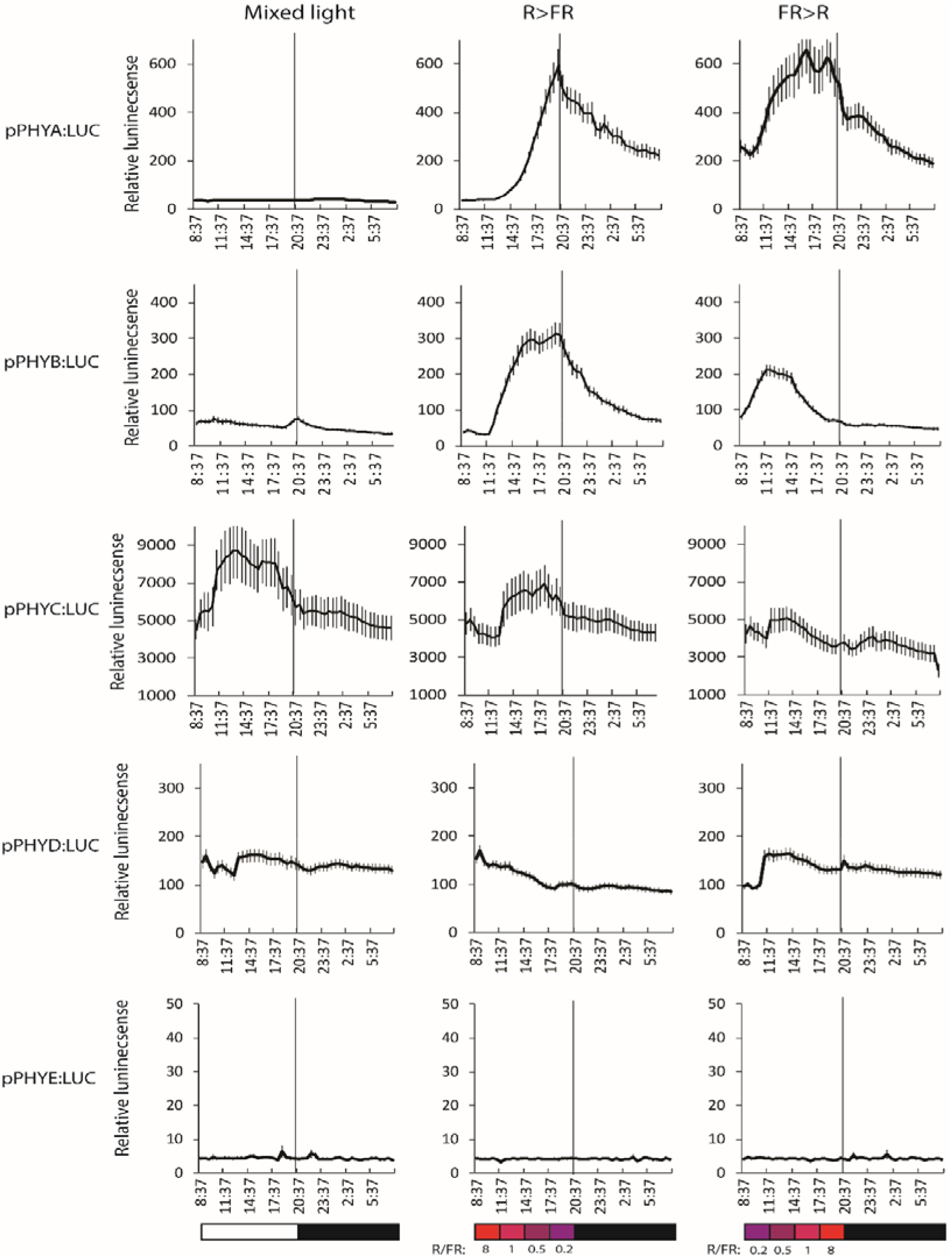
pPHY:LUC activity in WT rosette plants in response to changing R:FR ratios. pPHY:LUC in WT plants were grown in growth cabinets under diurnal fluorescent WL (12L/12D) for 25 days. Reporter plants were sprayed with substrate luciferin (1 mM) and one day later placed in LUMINATOR for adjustment under diurnal mixed R+B+FR for one day. Subsequently rosette plants were exposed to R light with increasing levels of FR (in blocks of 3 hours), resulting in R:FR ratios of 8, 1, 0.5 and 0.2. After the following night plants were exposed to the reverse light regime. Luciferin (1 mM) solution was sprayed once per day. LUC activity images were obtained every half hour (7 min. exposure) for each full diurnal cycle. The relative LUC activity was quantified in ImageJ and corrected for background signal. At least 7 replicate rosette plants were used for each reporter line. Error bars represent mean ±SE. The vertical line indicates the day to night transition.

When light treatment goes from deep shade to mild shade, only pPHYB:LUC shows a direct upregulation under R:FR=0,2, levels off at R:FR=0,5 and subsequently decreases under R:FR=1 and 8. While pPHYA:LUC shows the strongest response to R:FR=0,2 at end of day, when this shade condition is given at the start of the day pPHYA:LUC shows no response and only increases in activity at R:FR=0,5 or higher (Fig. 5). In general pPHYC:LUC decreases when R:FR increases, while in general pPHYD:LUC shows the opposite response and shows decrease in activity when R:FR increases. Overall, the results indicated changes in pPHY:LUC activity as function of shade, but dependent on time of day shade conditions are given. Subsequently the role of individual phytochromes is in the shade response of the different pPHY:LUC reporters was determined.

### PHYE is required for pPHYA:LUC response to different levels of FR added to R

To determine the role of each of the five phytochromes in the response to different R:FR ratios, pPHY:LUC reporter activities were quantified in all phytochrome mutant backgrounds (Fig. S2A-E). Here only the big effects on PHYA and PHYB expression are discussed. Expression of pPHYA:LUC is mostly affected by PHYD and PHYE: expression of pPHYA:LUC is much increased in the *phyD* mutant, while the upregulation of pPHYA:LUC under increasing FR is absent in the *phyE* mutant background (Fig. 6). This confirms a specific role for PHYE in FR light sensing in the regulation of PHYA gene expression. In contrast, the classical sensor for FR responses PHYA had only a small effect on FR-induction of PHYA-LUC (Fig. S2). The pPHYB:LUC activity is mostly affected by PHYB itself (Fig. S2B). The effect of PHYC, PHYD and PHYE on PHYB expression is conditional: they have little effect on pPHYB:LUC activity when going from high R:FR to low R:FR, but these phytochromes act as suppressor of pPHYB:LUC expression when light changes from low R:FR to high R:FR(Fig. S2B).

**Fig.6.**
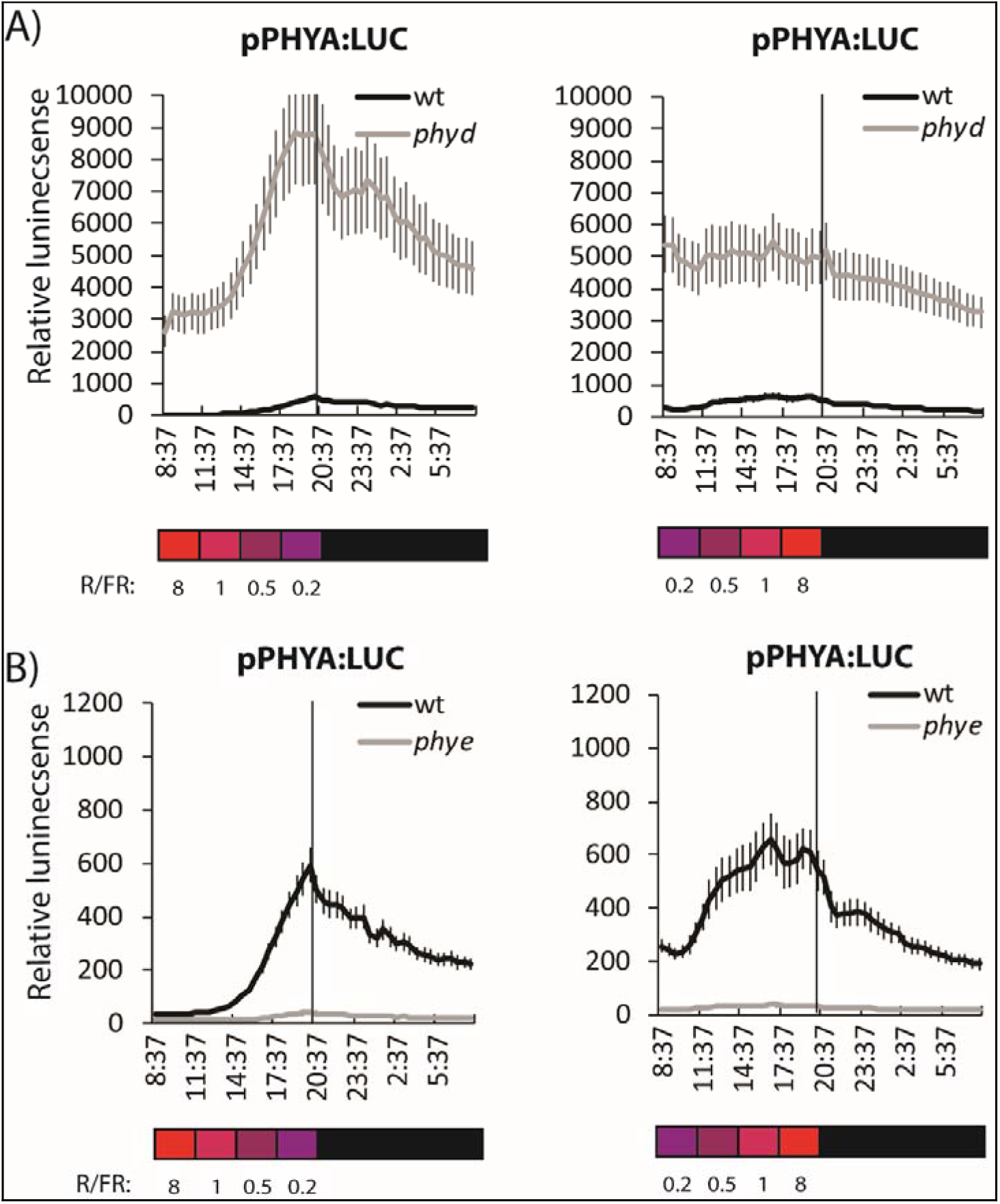
pPHYA:LUC activity in WT and *phyD* and *phyE* mutant in response to changing R:FR ratios. pPHYA:LUC in WT, *phyD* and *phyE* plants were grown in growth cabinets under diurnal fluorescent WL (12L/12D) for 25 days one day later placed in LUMINATOR for adjustment under diurnal mixed R+B+FR for one day. Subsequently rosette plants were exposed to R light with increasing levels of FR (in blocks of 3 hours), resulting in R:FR ratio’s of 8,1, 0.5 and 0.2. After the following night plants were exposed to the reverse light regime. Every day once at 11AM plants were sprayed with substrate luciferin (1 mM) solution. LUC activity images were obtained every half hour (7 min. exposure) for each full diurnal cycle. The relative LUC activity was quantified in ImageJ and corrected for background signal. Number of replicate seedlings for each reporter line: N=7. Error bars represent mean ±SE. A: pPHYA-LUC activity in WT, *phyD.* B: pPHYA-LUC activity in WT and *phyE.* Note that for activity in *phyD* mutant the scale of relative LUC activity was adjusted. Black: PHYA-LUC in WT, grey: PHYA-LUC in *phy* mutant. (Expression of all pPHY-LUC reporter lines in WT and pŕψ-mutants under changing R:FR is given in Fig. S4).

Results for the PHY:LUC reporters reflect the activity of endogenous transcription factors on the inserted transgenes. To determine if the reporter activity also reflecst activity of the endogenous PHY genes some of the strong effects of PHYD and PHYE on expression of PHYA, strong effect of PHYE on expression of PHYB were validated by qPCR. Results shows a different dynamics of induction of endogenous PHYA under FR, indicating that the normal context of the PHYA gene is far more buffered against effects of FR than the pPHYA:LUC reporter. However, endogenous PHYA is induced after 10 hr of FR and this induction is absent in the *phyE* mutant, confirming the interaction as shown for the pPHYA:LUC reporter. Moreover, also the strong suppression by PHYD of PHYA expression is confirmed for the endogenous PHYA gene (Fig. 7A). The endogenous PHYB gene shows rapidly upregulation under FR and this upregulation is reduced in the *phyE* mutant for the early response peak, confirming a role for PHYE in FR responses for PHYB expression (Fig. 7B).

**Fig.7.**
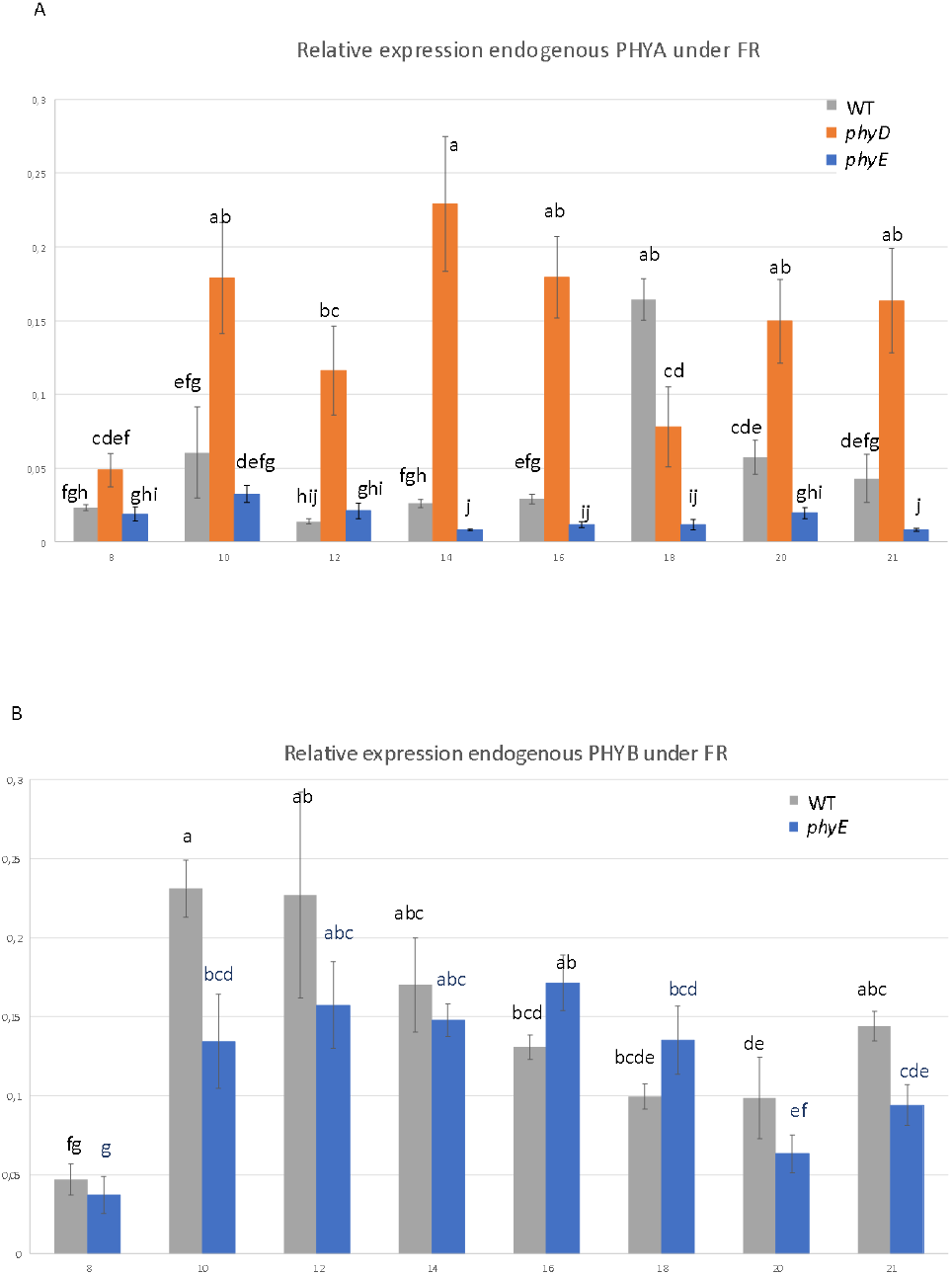
qPCR validation for endogenous PHYA and PHYB expression under FR. Plants were exposed to the same cFr as used in Figure 3. A: Endogenous. PHYA mRNA levels in WT, *phyD* and *phyE* mutant backgrounds were quantified. B: Endogenous PHYB mRNA levels in WT and *phyE* mutant background were quantified. Error bars with different letters indicate statistical significance difference at the P<0.01 level analysed by two-way ANOVA with LSD *post hoc* test.

## Discussion

### Mutual transcriptional regulation also at the basis of complex interaction between PHYs

The role of the different phytochrome gene family members of the Arabidopsis has mostly been studied for germination, hypocotyl elongation and flowering responses and these phenotypes have also been used to dissect the complex genetic interactions between the different Arabidopsis phytochrome genes (Sanchez-Lamas, Lorenzo et al. 2016). Here we have revealed a potential basis of these complex interactions by revealing that phytochrome genes influence each other’s transcription. The interactions are complex because they may depend on given light conditions. For instance, the activity of PHYB on its own expression reverses depending on the light conditions: under mixed light and R LED PHYB suppresses its own expression, while under FR PHYB is required for the full induction response (Fig. 5). This light dependent activity is also visible in the experiment with different R:FR light treatments, which shows that PHYB strongly suppresses its own gene expression under mixed LED light, but is required for the response to R plus added FR light (Fig. S2B).

The transcriptional factors that act on the pPHY:LUC reporters show a strong response in some of the single phytochrome mutants (Figs. 2, 4, 5). However, validation of the pPHYA:LUC and pPHYB:LUC reporter responses also indicates that endogenous PHYA and PHYB gene expression may be more buffered against the effect of single phytochrome mutation (Fig. 7). Nevertheless, major effects revealed by the pPHY:LUC reporters were validated for endogenous PHYA and PHYB (Fig. 7): PHYE is required for full FR induced transcription of PHYA and PHYB and PHYD represses PHYA transcription.

### PHYD is a constitutive suppressor of PHYA gene transcription

One of the strongest and consistent interactions these studies have uncovered is the suppression of PHYA transcription by PHYD at different stages of development and under different light conditions (Figs 2,6,7). Assuming that this interaction is cell specific, this implies that PHYD is expressed in most cells that express PHYA. The biological function of this suppression of PHYA by PHYD is at present not clear. Our results are consistent with the complementary expression profiles of PHYA and PHYD in developing and dry seeds (low PHYA, high PHYD), and imbibed seeds (high PHYA and low PHYD) (Toufighi, Brady et al. 2005). The function of PHYD thus could to be to limit PHYA expression in developing seeds. PHYD can form a homodimer and heterodimers with PHYB, PHYC and PHYE (Sharrock and Clack 2004). None of the mutants of *phyB, phyC* or *phyE* show a strong effect on pPHYA:LUC expression (Fig. 2), suggesting that it may be the combined loss of PHYD homodimers and heterodimers that are responsible for the strong upregulation of PHYA expression in the *phyD* mutant. Future analysis will have to show how PHYA expression is affected in double and triple PHY mutants. The higher expression level of PHYA in the *phyD* mutant background may relate to the different phenotypes that have been described for the Arabidopsis *phyD* mutant (Christians, Gingerich et al. 2012, Sanchez-Lamas, Lorenzo et al. 2016).

### Gating of the FR-induced PHY transcriptional response

The strong induction of both PHYA and PHYB is not only an artifact of the unnatural pure FR light condition (Figs 3 and 4), but is also seen at more physiological levels of R:FR (Figs S4, 5 and 6). The responses to different ratios of R:FR are dependent on time of day at which they are given, with the strongest response near end of day and the strongest suppression of a FR-induced transcriptional response at the start of the day. Overall, this suggests a gating of the FR transcriptional response of PHYA and PHYB by the circadian clock and this gating coincides in gating of the rapid FR elongation response observed in seedlings (Salter, Franklin et al. 2003). Expression of phytochrome genes is seedlings is strongly regulated by the clock (Toth, Kevei et al. 2001). However, the amplitude in PHY expression by clock regulation seems to be very much reduced in mature rosetted plants (Figs 3 and 5). The reciprocal interaction between PHYs and the clock on PHY gene transcription, combined with the interactions between different PHYs on PHY gene transcription as function of light condition, may explain the different phases for the FR light responses for PHYA and PHYB.

### Potential consequences for the total phytochrome protein pool and heterodimer formation

Phytochrome activity under different light conditions has mainly been studied for signalling downstream of PHY^pfr^, which is both a function of the total phytochrome protein pool and the equilibrium between active PHY^pfr^ and inactive PHY^Pr^. Although it is well established that the fraction of activated PHY protein is determined by R:FR ratio, here we have shown that under specific light conditions also the input of PHY protein may change. The increased transcription of PHYA and PHYB in response to FR may increase the total PHYA and PHYB protein pool. Potentially, this may compensate a little for the shift in the equilibrium between PHY^pfr^ and PHY^pr^ under FR towards inactive PHY^pr^. In addition, an increased PHYA and PHYB protein pool in response to FR increases the potential for light signaling when conditions favour activation of PHY protein again. This could have potential applications in manipulating PHY action in greenhouses to steer crop performances.

Phytochrome interactions studies have revealed that PHYC may form heterodimer with PHYB and PHYD and that PHYC may not exist as homodimer (Clack, Shokry et al. 2009). The relative high expression of PHYC compared to that of PHYB, PHYD and especially PHYE (which is expressed at very low levels), suggest that PHYB and PHYD may preferentially exist as heterodimer with PHYC and that PHYB/D and PHYB/E heterodimers are only formed as minor components. Removal of PHYC from this pool of interacting phytochromes could therefore result in a substantial increase in the pool of PHYB/D and PHYB/E heterodimers. The induction of PHYB expression under FR is strongly affected by PHYB and PHYE, but not PHYC or PHYD. This could suggest that the induction of PHYB expression under FR may be mostly through PHYB/E heterodimers (Hofmann 2009).

### Transcriptional interaction between PHY genes in dicots and monocots

The classical high irradiance response (HIR) of Arabidopis is characterized by the suppression of hypocotyl elongation. Both PHYA and PHYB are involved in this HIR response (Quail, Boylan et al. 1995), but PHYB is mostly responsible for HIR under cR light (R-HIR) (Nagatani, Kay et al. 1991, Reed, Nagpal et al. 1993) and PHYA predominantly for the HIR responses under cFR light (FR-HIR) (Hartmann 1967, Nagatani, Reed et al. 1993, Parks and Quail 1993, Whitelam, Johnson et al. 1993, Casal, Candia et al. 2014, Possart, Fleck et al. 2014). The strong induction of PHYA promoter activity under FR light may be considered as a novel FR-HIR response. However, not PHYA but PHYE is involved in this FR-HIR induction of PHYA gene activity (Figs 6, 7). Phytochromes are classified as either Type I, which are activated by far-red light, or Type II that are activated by red light (Li, Li et al. 2011), although phytochrome Type I and Type II may also be defined by the phytochrome protein stability in light. For Arabidopsis only PHYA has been classified as a Type I phytochrome, as it is responsible for many FR light induced responses and is instable in the light. With the extension of FR-HIR responses beyond seedling de-etiolation to PHY gene expression under FR, the classification of Arabidopsis PHYE as type II phytochrome may need reconsideration.

Contrary to the five PHY genes in Arabidopsis, rice has only a PHYA, PHYB, and PHYC. Presumably, the PHYA, PHYB and PHYC were already formed before the formation of gymnosperms, as both monocotyledons and dicotyledons contain representatives of PHYA, PHYB, and PHYC. In dicotyledonous plants, duplications of the PHYB progenitors resulted in the PHYE subfamily and, specifically in Arabidopsis, another duplication event of PHYB resulted in PHYD (Clack, Mathews et al. 1994). In contrast, grasses lack the PHYD and PHYE members of the PHYB subfamily. While the PHYC in Arabidopsis is a type II phytochrome, in rice, PHYC mediates FR-HIR de-etiolation and therefore could be considered a Type I phytochrome (Takano, Inagaki et al. 2005). Future research will have to show whether or how the effect of individual PHY genes in dicots and monocots relates to mutual transcriptional regulation of PHY gene expression and how this modulates phenotypic responses to different light conditions.

## Acknowledgement

This research was in part funded by the STW Project (13149) ‘Compact Plants’.

## Supplemental Figure legends

**Fig.S1.**
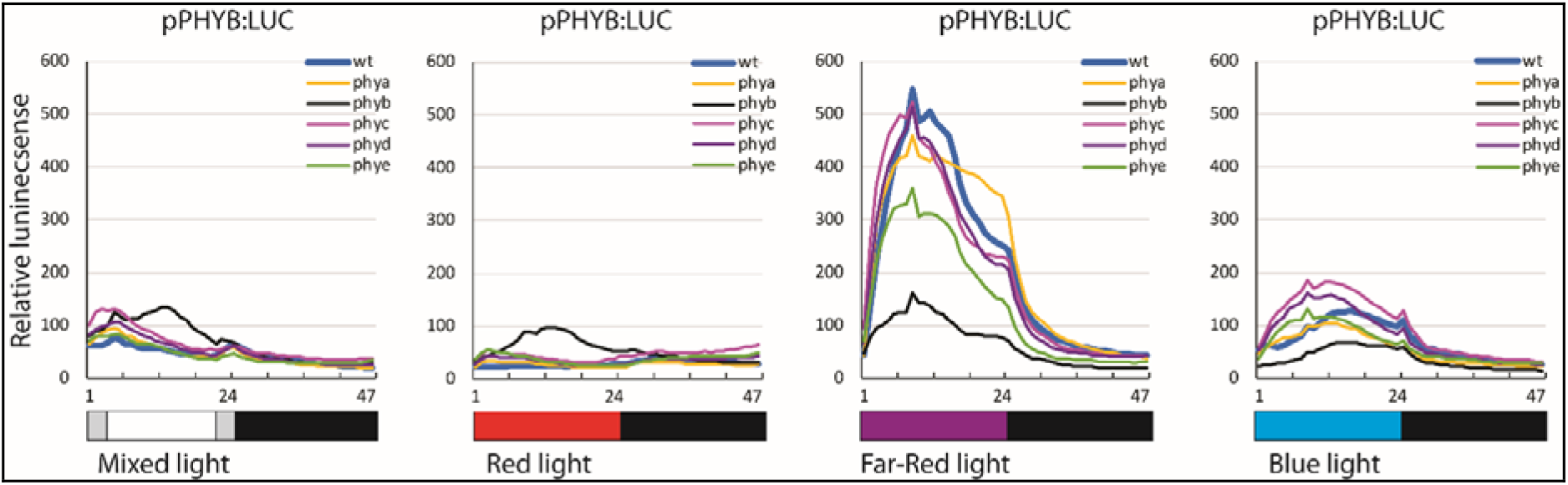
Diurnal pPHYB:LUC activity in WT and *phy* mutant plants under mixed R+B+FR light. Seeds of pPHYB-LUC reporter in WT and the five single PHY mutant backgrounds were stratified and germinated in growth cabinets under diurnal fluorescent WL (12L/12D). At 14 days after germination seedlings were sprayed with substrate luciferin (1 mM) and one day later placed in LUMINATOR for adjustment under diurnal R+B+FR for one day. Subsequently plants were exposed to light regimes 12mixed/12D, 12R/12D, followed by 12FR/D and finally 12B/12D. Luciferin (1 mM) solution was sprayed once per day. LUC activity images were obtained every half hour (7 min. exposure) for each full diurnal cycle. The relative LUC activity was quantified in Image J and corrected for background signal. Number of replicate seedlings for each reporter line: N=6. Error bars represent mean ±SE.

**Fig.S2.**
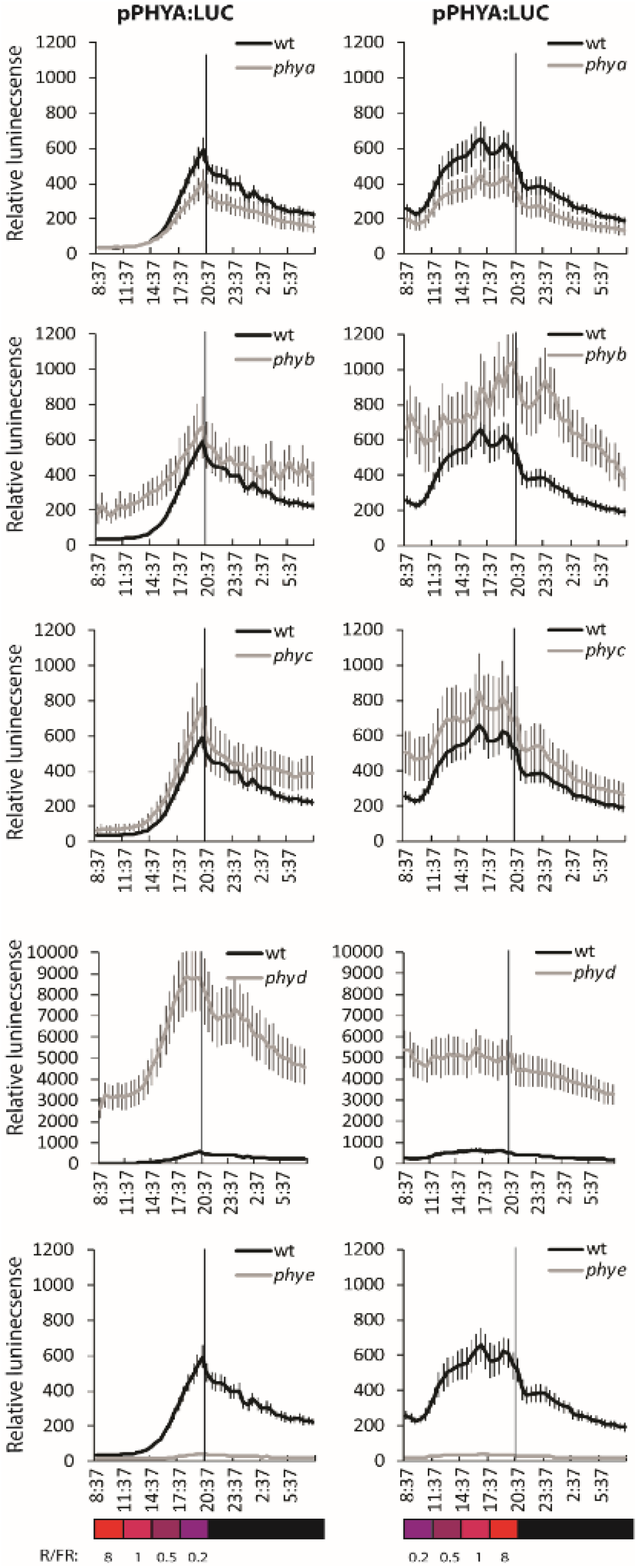
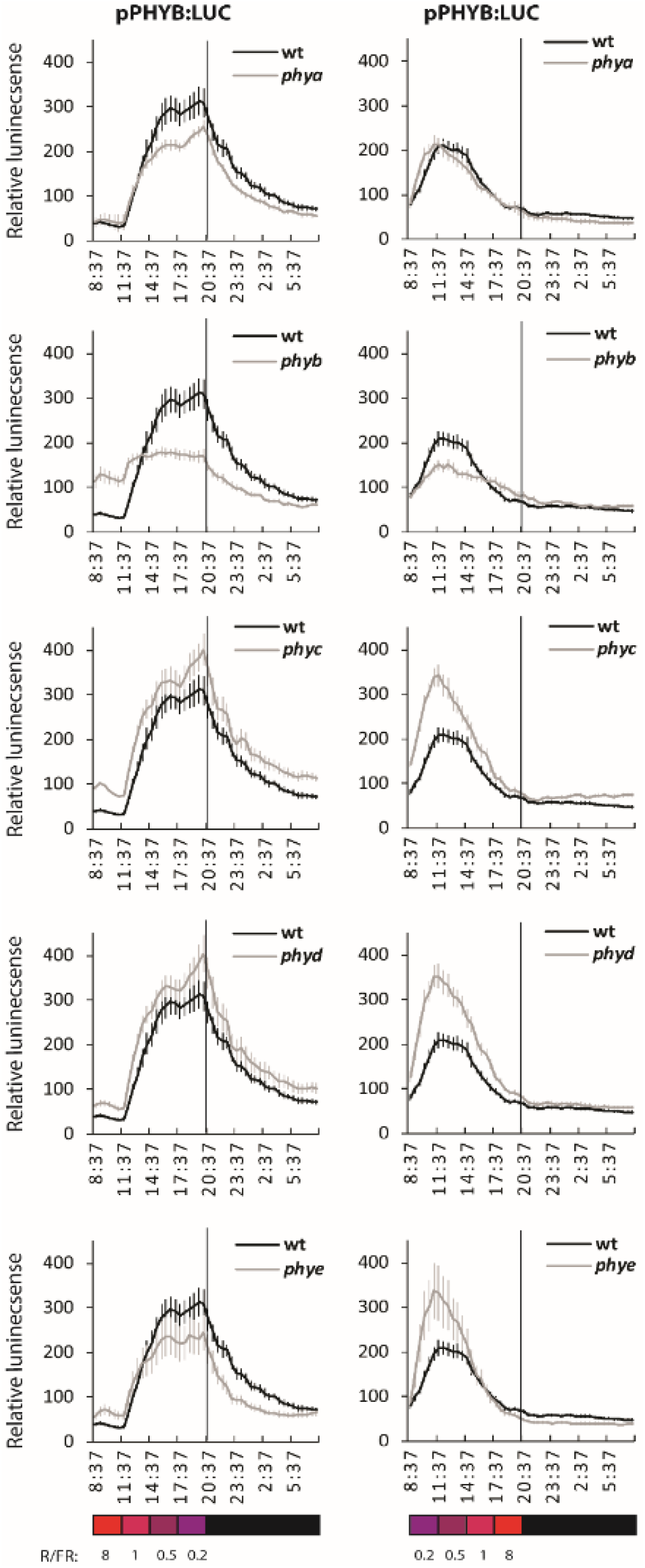
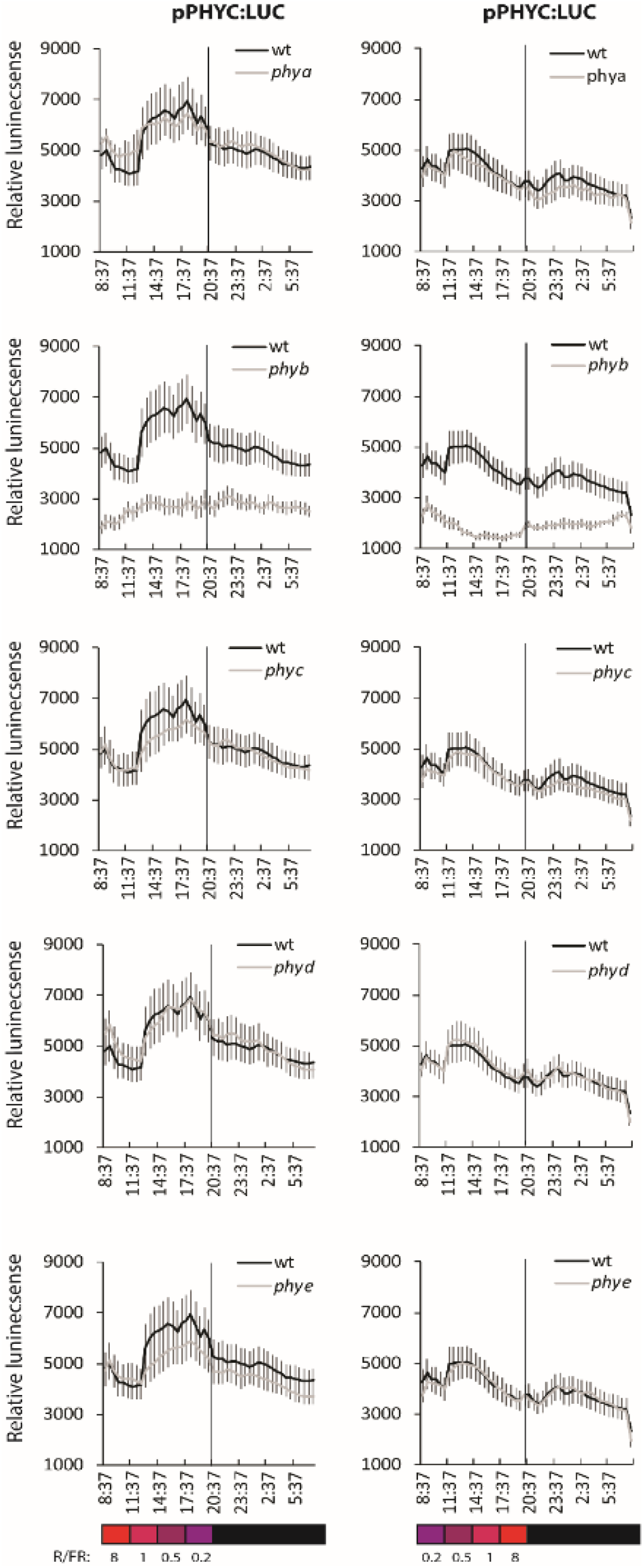
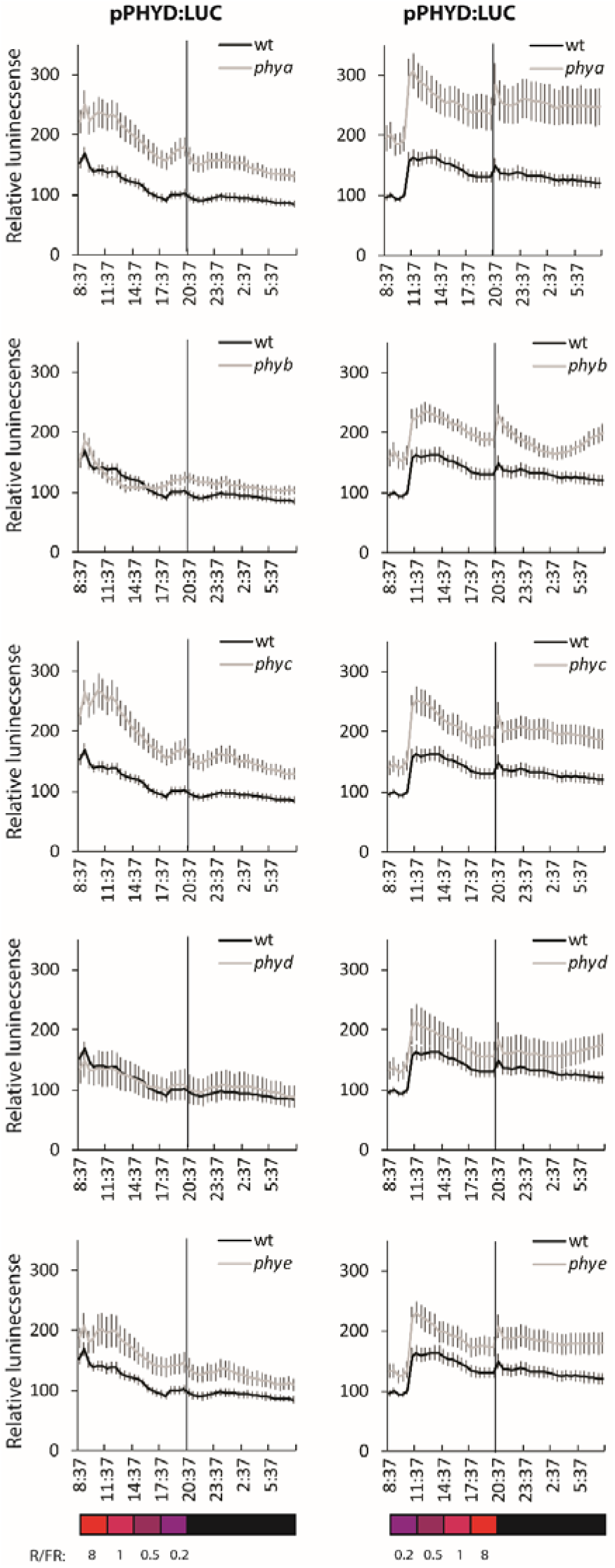
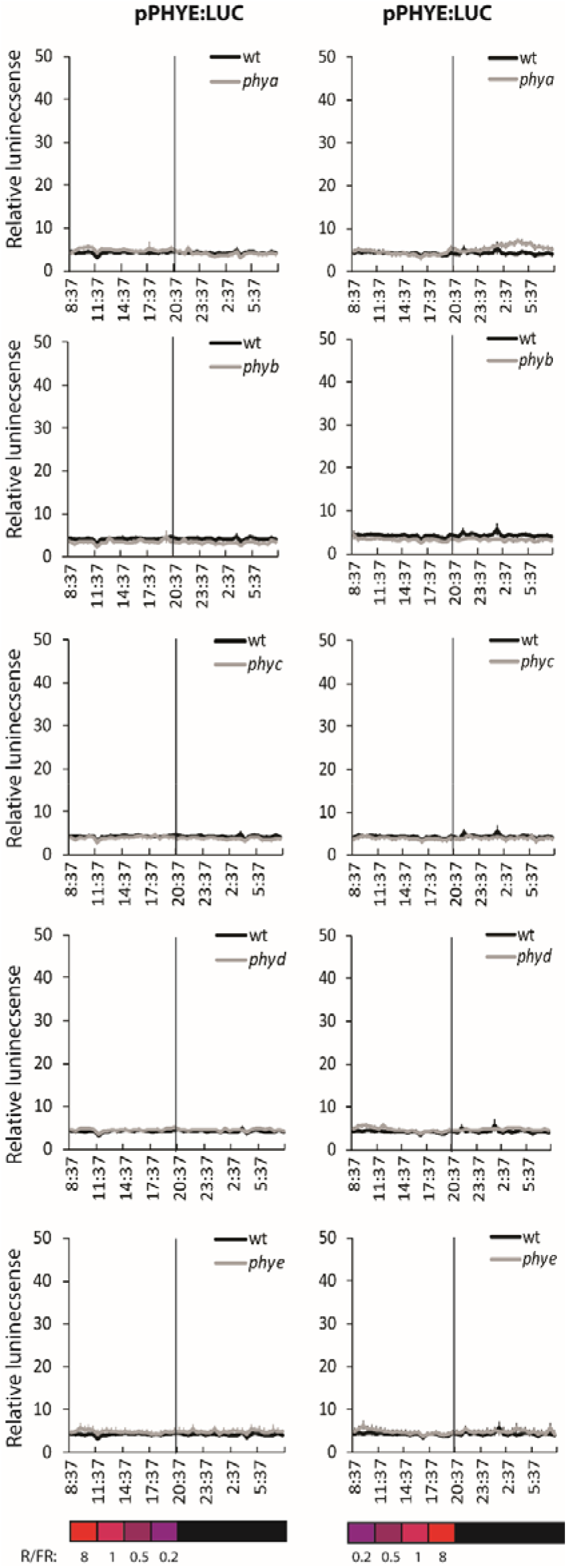
A-E. pPHY:LUC activity in *phy* mutant compared WT rosette plants in response to changing R:FR ratios. Plants were grown under diurnal fluorescent WL (12L/12D). At 25 days after germination plants were sprayed with substrate luciferin (1 mM) and one day later placed in LUMINATOR for adjustment under diurnal mixed R+B+FR for one day. Subsequently plants were exposed to a fixed level of R with increasing levels of FR (in blocks of 3 hours), resulting in R:FR ratio’s of 8, 1,0.5 and 0.2. After the following night plants were exposed to the reverse light regime. Luciferin (1 mM) solution was sprayed once per day. LUC activity images were obtained every half hour (7 min. exposure) for each full diurnal cycle. The relative LUC activity was quantified in Image J and corrected for background signal. Number of replicate seedlings for each reporter line: N=6. Error bars represent mean ±SE. A: pPHYA:LUC activity in WT and the five *phy* mutant backgrounds. Note that for the *phyD* mutant background the scale of relative LUC activity is different. B: pPHYB:LUC activity in WT and the five *phy* mutant backgrounds. C: pPHYC:LUC activity in WT and the five *phy* mutant backgrounds. D: pPHYD:LUC activity in WT and the five *phy* mutant backgrounds. E: pPHYE:LUC activity in WT and the five *phy* mutant backgrounds. Note that activity is barely above background.

## Supplemental Tables

**Table-S1.**
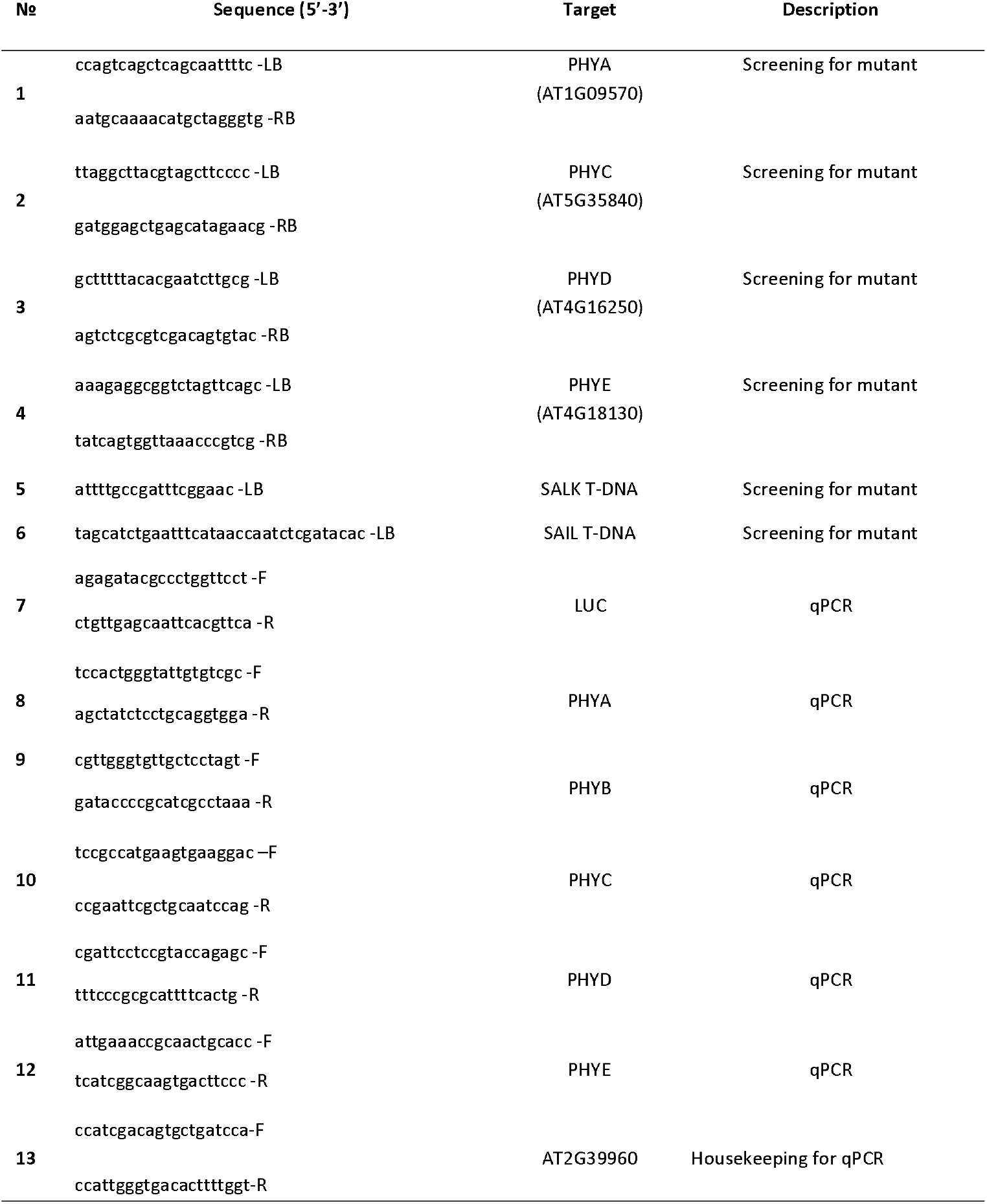
Primers used in this study.

**Table-S2.** List of reporter lines were used in study. The pPHY:LUC reporter was created in Col-0 and its crossed with different mutant background lines.

| LUC reporter lines, constructs | Description | Plasmid notes | Ref |
| --- | --- | --- | --- |
| pPHYA:LUC<sup>WT</sup> | pPHYA:LUC reporter in Col-0 | *Hyg<sup>R</sup>*/pPCVH | In study |
| pPHYA:LUC<sup>phyA</sup> | pPHYA:LUC reporter crossed to *phyA* mutant | *Hyg<sup>R</sup>*/pPCVH | In study |
| pPHYA:LUC<sup>phyB-9</sup> | pPHYA:LUC reporter crossed to *phyB-9* mutant | *Hyg<sup>R</sup>*/pPCVH | In study |
| pPHYA:LUC<sup>phyC</sup> | pPHYA:LUC reporter crossed to *phyC* mutant | *Hyg<sup>R</sup>*/pPCVH | In study |
| pPHYA:LUC<sup>phyD</sup> | pPHYA:LUC reporter crossed to *phyD* mutant | *Hyg<sup>R</sup>*/pPCVH | In study |
| pPHYA:LUC<sup>phyE</sup> | pPHYA:LUC reporter crossed to *phyE* mutant | *Hyg<sup>R</sup>*/pPCVH | In study |
| pPHYB:LUC<sup>WT</sup> | pPHYB:LUC reporter in Col-0 | *Hyg<sup>R</sup>*/pPCVH | In study |
| pPHYB:LUC<sup>phyA</sup> | pPHYB:LUC reporter crossed to *phyA* mutant | *Hyg<sup>R</sup>*/pPCVH | In study |
| pPHYB:LUC<sup>phyB-9</sup> | pPHYB:LUC reporter crossed to *phyB-9* mutant | *Hyg<sup>R</sup>*/pPCVH | In study |
| pPHYB:LUC<sup>phyC</sup> | pPHYB:LUC reporter crossed to *phyC* mutant | *Hyg<sup>R</sup>*/pPCVH | In study |
| pPHYB:LUC<sup>phyD</sup> | pPHYB:LUC reporter crossed to *phyD* mutant | *Hyg<sup>R</sup>*/pPCVH | In study |
| pPHYB:LUC<sup>phyE</sup> | pPHYB:LUC reporter crossed to *phyE* mutant | *Hyg<sup>R</sup>*/pPCVH | In study |
| pPHYC:LUC<sup>WT</sup> | pPHYC:LUC reporter in Col-0 | *Hyg<sup>R</sup>*/pPCVH | In study |
| pPHYC:LUC<sup>phyA</sup> | pPHYC:LUC reporter crossed to *phyA* mutant | *Hyg<sup>R</sup>*/pPCVH | In study |
| pPHYC:LUC<sup>phyB-9</sup> | pPHYC:LUC reporter crossed to *phyB-9* mutant | *Hyg<sup>R</sup>*/pPCVH | In study |
| pPHYC:LUC<sup>phyC</sup> | pPHYC:LUC reporter crossed to *phyC* mutant | *Hyg<sup>R</sup>*/pPCVH | In study |
| pPHYC:LUC<sup>phyD</sup> | pPHYC:LUC reporter crossed to *phyD* mutant | *Hyg<sup>R</sup>*/pPCVH | In study |
| pPHYC:LUC<sup>phyE</sup> | pPHYC:LUC reporter crossed to *phyE* mutant | *Hyg<sup>R</sup>*/pPCVH | In study |
| pPHYD:LUC<sup>WT</sup> | pPHYD:LUC reporter in Col-0 | *Hyg<sup>R</sup>*/pPCVH | In study |
| pPHYD:LUC<sup>phyA</sup> | pPHYD:LUC reporter crossed to *phyA* mutant | *Hyg<sup>R</sup>*/pPCVH | In study |
| pPHYD:LUC<sup>phyB-9</sup> | pPHYD:LUC reporter crossed to *phyB-9* mutant | *Hyg<sup>R</sup>*/pPCVH | In study |
| pPHYD:LUC<sup>phyC</sup> | pPHYD:LUC reporter crossed to *phyC* mutant | *Hyg<sup>R</sup>*/pPCVH | In study |
| pPHYD:LUC<sup>phyD</sup> | pPHYD:LUC reporter crossed to *phyD* mutant | *Hyg<sup>R</sup>*/pPCVH | In study |
| pPHYD:LUC<sup>phyE</sup> | pPHYD:LUC reporter crossed to *phyE* mutant | *Hyg<sup>R</sup>*/pPCVH | In study |
| pPHYE:LUC<sup>WT</sup> | pPHYE:LUC reporter in Col-0 | *Hyg<sup>R</sup>*/pPCVH | In study |
| pPHYE:LUC<sup>phyA</sup> | pPHYE:LUC reporter crossed to *phyA* mutant | *Hyg<sup>R</sup>*/pPCVH | In study |
| pPHYE:LUC<sup>phyB-9</sup> | pPHYE:LUC reporter crossed to *phyB-9* mutant | *Hyg<sup>R</sup>*/pPCVH | In study |
| pPHYE:LUC<sup>phyC</sup> | pPHYE:LUC reporter crossed to *phyC* mutant | *Hyg<sup>R</sup>*/pPCVH | In study |
| pPHYE:LUC<sup>phyD</sup> | pPHYE:LUC reporter crossed to *phyD* mutant | *Hyg<sup>R</sup>*/pPCVH | In study |
| pPHYE:LUC<sup>phyE</sup> | pPHYE:LUC reporter crossed to *phyE* mutant | *Hyg<sup>R</sup>*/pPCVH | In study |

